# Relationship Between Circulating FGF21 and Physiological and Lifestyle Characteristics, an Exploratory Study

**DOI:** 10.1101/2024.12.26.630421

**Authors:** Matthew Peterson, Kathleen Adair, LesLee Funderburk

## Abstract

Fibroblast growth factor 21 (FGF21) is a biomarker that has been linked to metabolic health. This study was conducted to examine the relationship between FGF21 and its purported upstream regulators and downstream targets in healthy humans. Male and female participants completed three study visits. During visit 1 anthropometrics were measured and a VO_2peak_ test was conducted on a stationary bike. During visits 2 and 3 resting metabolic rate, heart rate, and blood pressure were recorded and a blood sample was taken. Results from visits 2 and 3 were averaged before statistical analysis. Additionally, participants completed a food frequency questionnaire to detail typical diet over the previous month and had three days of measured physical activity and sleep. Moderate correlations between FGF21 and measured sleep (*r* = 0.34, *p* = 0.05), saturated fat intake (*r* = −0.37, *p* = 0.04), and blood urea nitrogen levels (*r* = −0.47, *p* = 0.01) were observed. Additionally, greater intensities of daily physical activity were associated with lower FGF21 levels. Serum FGF21 levels may be influenced by lifestyle variables such as sleep and physical activity, diet, and other circulating biomarkers.

**Highlights:**

- FGF21 is positively correlated with sleep in healthy humans.
- Sugar intake is positively correlated with FGF21 concentrations in humans.
- Blood urea nitrogen concentrations are negatively correlated with FGF21 in humans.

## Introduction

Fibroblast growth factor 21 (FGF21) is a member for the FGF19 subfamily of FGF molecules.(Hu *et al*., 2013, p. 23) The FGF19 subfamily is unique in that its members circulate throughout the body.(Goetz *et al*., 2007) As a result, FGF21 has a broad range of endocrine-like effects on the body. Studies in mice have demonstrated that pharmacological administration of FGF21 is able to promote weight loss, reverse type II diabetes, and protect against the detrimental effects of a high fat diet.(Ding *et al*., 2012) This has led to speculation that endogenous levels of FGF21 may be an important biomarker for predicting metabolic disease.

The study of FGF21 in humans has been complicated by the fact that FGF21 is differentially regulated in humans versus mice (Fazeli *et al*., 2015) and a large variance in resting FGF21 concentrations has also been observed, even among the same individuals.(Gälman *et al*., 2008; Christodoulides *et al*., 2009) Additionally, the upstream regulators and downstream effectors in the FGF21 pathway are multifactorial and oftentimes exhibit paradoxical relationships.(Uebanso *et al*., 2011)

One of the first paradoxes witnessed in the regulation of FGF21 is with obesity. Observational studies have indicated that obese individuals have greater levels of circulating FGF21 than do normal weight individuals.(Zhang *et al*., 2008) Similar to weight status, lifestyle variables such as physical activity and dietary intake are associated with circulating levels of FGF21. Acute bouts of exercise have been shown to increase FGF21 levels post-exercise.(Kim *et al*., 2013; Hansen *et al*., 2015; Morville *et al*., 2018) However, this increase is transient.(Hecht *et al*., 2012) Studies investigating the effect of habitual exercise have similarly shown a positive relationship between physical activity and FGF21.(Cuevas-Ramos *et al*., 2010) Transcription of the *FGF21* gene can be promoted by both carbohydrates through the carbohydrate-response element-binding protein (ChREBP)(Uebanso *et al*., 2011) and by fatty acids through the peroxisome proliferator-activated receptor alpha (PPARα) transcription factor.(Christodoulides *et al*., 2009; Uebanso *et al*., 2011) This further underscores the paradox of FGF21 regulation, as it can be regulated through both feeding and fasting mechanisms. Additionally, nutrients that are metabolized in the liver have been shown to have acute effects on circulating FGF21 levels.(Dushay *et al*., 2015; Søberg *et al*., 2018) Complicating matters, not only does FGF21 respond to nutrient intake, but there is evidence to suggest that FGF21 may also influence nutrient intake and preference through its actions in the central nervous system.(Adams and Gimeno, 2016; Talukdar *et al*., 2016)

In addition to lifestyle factors, FGF21 may also be associated with other circulating factors. Prior research has demonstrated a positive correlation between blood glucose and FGF21 levels.(Cuevas-Ramos *et al*., 2010; Gómez-Ambrosi *et al*., 2017) Research in animals has shown a relationship between FGF21, fluid balance and kidney function,(Turner *et al*., 2018) which might mean FGF21 is associated with human markers of kidney function as well as the regulation of blood pressure. While FGF21 may be expressed in a variety of tissues, the majority of evidence in humans suggests that under normal conditions circulating FGF21 is derived from the liver;(Hansen *et al*., 2015) however, there has been minimal work examining the relationship between FGF21 and other proteins produced by the liver. Finally, one of the major findings from a study that investigated the pharmacological dosing of an FGF21 analog was an improvement in dyslipidemia,(Gaich *et al*., 2013) indicating that FGF21 may promote the beneficial regulation of blood cholesterol (i.e. decreasing LDL and increasing HDL).

While much may be known about upstream and downstream pathways of FGF21, the extent to which these processes occur *in vivo* in healthy individuals are unknown. Therefore, the purpose of this study was to conduct an exploratory analysis on relationships between FGF21 and factors that are believed to either regulate FGF21 or serve as downstream targets of FGF21 action.

## Materials and Methods

### Experimental Design

This study was an observational study that was conducted as part of a larger investigation into the relationship between metabolism and FGF21. The study was approved by the local Institutional Review Board. The study required three visits. On visit one, anthropometric measurements and the VO_2peak_ test were conducted. Visits 2 and 3 were identical: Following an overnight fast, participants reported to the lab in the morning at a time that was consistent with their normal sleep schedule. Upon arriving to the lab, the participant’s body weight was recorded, and their resting metabolic rate was measured for fifteen minutes. During the first five minutes of the resting measures, a blood sample was obtained, and their blood pressure was measured. Participants were asked to match their diet and physical activity as closely as possible for the days prior to visit 2 and visit 3; as well as to avoid vigorous physical activity and ethanol consumption during this time. To account for the high variability in FGF21 measures, the results from visits 2 and 3 were averaged before conducting the statistical analyses.

### Participants

Thirty apparently healthy, physically active adults were recruited for this study. Fifteen participants were female and fifteen were male. All participants were briefed on study procedures and provided written informed consent prior to enrolling in the study. Recruitment criteria included: eighteen to forty-five years of age, participated in at least thirty minutes of moderate activity three days a week for the past three months or longer, normal weight as indicated by a BMI between 18.5 and 24.9, and free of any cardiovascular, metabolic, or renal diseases.

### Blood Analyses

Blood was drawn in the morning, following an overnight fast of at least ten hours. Serum was separated and stored at −80 ℃ until the analysis was conducted. Serum FGF21 was measured using a commercially available enzyme-linked immunosorbent assay (ELISA) (R&D Systems, DF2100). A metabolic panel and lipid panel were completed and reported by Clinical Pathology Laboratories (Waco, TX) using the Roche COBAS automated methodology and the Roche Cobas enzymatic colorimetric generation 2, respectively.

### Anthropometrics

Height and weight were measured to the nearest 0.1 cm and 0.1 lbs, respectively, using a stadiometer and calibrated digital scale (Seca; Chino, CA). Both measurements were performed with the participant in exercise clothes, without shoes, but in stocking feet. Waist circumference was measured with a Gulick tape at the level of the umbilicus.

### Physiological Measurements

During the first visit, a VO_2peak_ test was conducted on a mechanically braked bike (Lode; Groningen, Netherlands), oxygen consumption was measured using a Parvo Medics TrueOne 2400 (Parvo Medics; Salt Lake City, UT). The participants maintained a pedaling cadence of 60rpm throughout the test. The bike resistance started at 50W and was increased 50W every two minutes. The test terminated when the participant either reached volitional fatigue or was no longer able to maintain a cadence of 60rpm for a period of thirty consecutive seconds (whichever came first). A VO_2peak_ was confirmed by a respiratory exchange ratio (RER) above 1.1 and plateau in VO_2_.

During the second and third visits, resting metabolic rate (RMR) was measured using the same Parvo Medic measuring system. RMR was measured with participants resting in a supine position for fifteen minutes, with the final five minutes used for analysis. Blood pressure was measured with the participant in the supine position using an automated blood pressure cuff (ADC E-sphyg 2; Hauppauge, NY).

### Sleep and Physical Activity

Participants were given an activity monitor (SenseWear by Bodymedia; Pittsburgh, PA) that measures sleep and physical activity. The participants were asked to wear the monitor for three days (one weekend day and two weekdays) outside of the study activities that represented their normal habits. The monitor had to be worn for at least twenty-three hours for the day to be considered valid.

### Diet

To assess the impact of typical diet on circulating FGF21 levels, participants were asked to fill out a food frequency questionnaire (DHQIII)(*DHQ3*, no date) that asked them about the types and quantities of food they typically consumed during the preceding month. The DHQIII provided information about the average macronutrient and micronutrient consumption of the participants over the previous month as well as calculated a healthy eating index (HEI) score.

### Statistics

FGF21 values were not normally distributed and were log-transformed prior to analysis. Correlations between FGF21 and variables related to either its downstream or upstream pathways were performed. A Pearson correlation was conducted for normally distributed data and a Spearman correlation was conducted for non-normally distributed data. There is some evidence to suggest that FGF21 may be differentially regulated based on biological sex,(Owen *et al*., 2013) therefore we present our results for the entire group and for each sex individually. Missing data were eliminated using listwise deletion. All analyses were conducted using SPSS version 26 (SPSS; Chicago, IL). An *a priori* alpha level of *p*<0.05 was adopted throughout. Supplemental tables and figures can be accessed at the following link: <u>10.6084/m9.figshare.19991213</u>.

## Results

### Participants

A total of 32 participants provided informed consent and were enrolled in this study. Two participants withdrew from the study due to time (n = 1) and lack of comfortability with the indirect calorimetry equipment (n = 1). Therefore, thirty participants completed this study (n = 15 females, n = 15 males); however, 5 had FGF21 levels that were below the minimum level of detection, providing a final sample of 25 participants (n = 13 females, n = 12 males). The average coefficient of variation for blood concentration of FGF21 between the two visits was 38.0% with a range of 1.1% to 105.0%. There were no statistically significant differences in serum FGF21 levels between males and females (*p>0.05*). A participant flow diagram is presented in supplemental figure 1 and demographic characteristics of our sample are shown in supplemental table 1.

### Age and Anthropometrics

For the whole sample, we observed a weak, positive correlation between age and FGF21 (*r* = 0.015, *p =* 0.40). Weight (*r =* −0.18, *p = 0.19*), BMI (*r =* −0.26, *p* = 0.10), and waist circumference (*r* = −0.15, *p* = 0.23) were all negatively associated with FGF21 levels. In females, a greater weight (*r* = −0.53, *p* =0.03), BMI (*r* = −0.49, *p* =0.04), and waist circumference (*r* = −0.53, *p* = 0.03) were all significantly associated with lower FGF21 levels. Results for the age and anthropometric analyses are shown in table 1.

**Table 1.** Correlations between FGF21 and anthropometric and activity variables. * = statistically significant correlation. BMI, body mass index (kg/m^2^); VO_2peak_, peak oxygen consumption; RMR, resting metabolic rate; RER, respiratory exchange ratio.

| Correlate | Whole Group |  | Females |  | Males |  |
| --- | --- | --- | --- | --- | --- | --- |
|  | *r* | *p* | *r* | *p* | *r* | *p* |
| Age | 0.05 | 0.40 | 0.22 | 0.23 | 0.07 | 0.41 |
| Weight | -0.18 | 0.19 | -0.53 | 0.03* | 0.43 | 0.08 |
| BMI | -0.26 | 0.10 | -0.49 | 0.04* | 0.20 | 0.27 |
| Waist Circum. | -0.15 | 0.23 | -0.53 | 0.03* | 0.37 | 0.20 |
| VO<sub>2peak</sub> | -0.26 | 0.11 | 0.19 | 0.27 | -0.71 | <0.01* |
| RMR | -0.20 | 0.17 | -0.48 | 0.04* | 0.30 | 0.17 |
| RER | -0.16 | 0.22 | -0.45 | 0.06 | 0.08 | 0.41 |
| Average daily activity | -0.11 | 0.30 | -0.20 | 0.26 | -0.02 | 0.48 |
| Sedentary time | 0.17 | 0.22 | 0.36 | 0.12 | -0.05 | 0.44 |
| Light activity | -0.15 | 0.24 | -0.2 | 0.47 | -0.23 | 0.24 |
| Moderate activity | -0.07 | 0.37 | -0.24 | 0.21 | 0.11 | 0.37 |
| Vigorous activity | -0.33 | 0.05 | 0.05 | 0.44 | -0.44 | 0.08 |
| Time in bed | 0.36 | 0.04* | 0.77 | <0.01* | -0.27 | 0.20 |
| Measured sleep | 0.34 | 0.05 | 0.71 | <0.01* | -0.26 | 0.21 |

### Metabolism, Activity, and Sleep

VO_2peak_ was negatively correlated with FGF21 (*r* = −0.26, *p* = 0.11) and in males this relationship was statistically significant with a strong effect size (*r* = −0.71, *p<*0.01). RMR was negatively related to FGF21 (*r* = −0.20, *p* = 0.17) and in females this was statistically significant with a strong effect size (*r* = −0.48, *p* = 0.04). Greater rates of FGF21 were associated with a lower RER (*r* = −0.16, *p* = 0.22) indicating that FGF21 was associated with greater amounts of fatty acid oxidation. Daily activity levels were associated with lower FGF21 levels (*r* = −0.11, *p* = 0.30), with a trend for greater intensity of activity being associated with lower FGF21 levels, while amount of sedentary time was positively associated with FGF21 (*r* = 0.17, *p* = 0.22). Sleep was positively correlated with FGF21 for the whole group (*r* = 0.34, *p* = 0.05) and in females this relationship was statistically significant and had a large effect size (*r* = 0.71, *p*<0.01); however, in males there was a negative correlation between FGF21 and sleep (*r* = −0.26, *p* = 0.21). Correlations for the metabolic, activity, and sleep data are shown in table 1.

### Diet

Serum FGF21 levels were compared to the average diet each participant consumed over the month prior to starting the study. Caloric consumption was weakly associated with circulating FGF21 levels (*r* = −0.11, *p* = 0.31). Fatty acid consumption was negatively related to blood FGF21 concentration (*r* = −0.23, *p* = 0.14), with all types of lipid species having a negative relationship; this effect was strongest for both saturated fat (*r* = −0.37, *p* = 0.04) and cholesterol (*r* = −0.42, *p* = 0.02). There was a positive relationship between carbohydrates and FGF21 (*r* = 0.12, *p* = 0.28) with all carbohydrate types sharing this positive relationship, the effect was strongest for sucrose (*r* = 0.31, *p* = 0.07). In females there was a statistically significant relationship for total sugar (*r* = 0.58, *p* = 0.03) and added sugar (*r* = 0.52, *p* = 0.04). Protein was negatively related to FGF21 (*r* = −0.29, *p* = 0.08). Ethanol was positively correlated with FGF21 (*r* = 0.20, *p* = 0.18). Finally, the healthy eating index (HEI) was positively associated with FGF21 for the whole group (*r* = 0.19, *p* = 0.19) and for females (*r* = 0.63, *p* = 0.01); however, in males, the HEI was negatively related to FGF21 levels (*r* = −0.20, *p* = 0.27). Dietary results are presented in table 2.

**Table 2.** Correlations between FGF21 and diet variables. * = statistically significant correlation. Kcal, kilocalories; HEI, healthy eating index.

| Correlate | Whole Group |  | Females |  | Males |  |
| --- | --- | --- | --- | --- | --- | --- |
|  | *r* | *p* | *r* | *p* | *r* | *p* |
| kcal | 0.11 | 0.31 | 0.04 | 0.45 | -0.23 | 0.24 |
| Fat | -0.23 | 0.14 | 0.11 | 0.36 | -0.43 | 0.08 |
| Saturated | -0.37 | 0.04* | -0.02 | 0.48 | -0.44 | 0.08 |
| Monounsaturated | -0.23 | 0.14 | 0.12 | 0.36 | -0.47 | 0.06 |
| Polyunsaturated | -0.09 | 0.35 | 0.28 | 0.19 | -0.32 | 0.16 |
| Cholesterol | -0.42 | 0.02* | -0.30 | 0.17 | -0.51 | 0.04 |
| Carbohydrates | 0.12 | 0.28 | 0.42 | 0.09 | -0.01 | 0.49 |
| Total Sugar | 0.21 | 0.16 | 0.58 | 0.03* | 0.05 | 0.44 |
| Added Sugar | 0.28 | 0.09 | 0.52 | 0.04* | 0.31 | 0.16 |
| Glucose | 0.13 | 0.27 | 0.43 | 0.08 | 0.09 | 0.39 |
| Sucrose | 0.31 | 0.07 | 0.46 | 0.07 | 0.16 | 0.31 |
| Fructose | 0.14 | 0.26 | 0.41 | 0.10 | 0.11 | 0.37 |
| Fiber | 0.12 | 0.29 | 0.25 | 0.22 | 0.08 | 0.41 |
| Protein | -0.29 | 0.08 | 0.09 | 0.39 | -0.32 | 0.15 |
| Dairy | -0.23 | 0.14 | 0.07 | 0.41 | -0.37 | 0.12 |
| Ethanol | 0.20 | 0.18 | 0.07 | 0.42 | 0.48 | 0.05 |
| HEI | 0.19 | 0.19 | 0.63 | 0.01* | -0.20 | 0.27 |

### Metabolic Panel

There was a weak, positive correlation between fasting blood glucose and FGF21 (*r* = 0.10, *p* = 0.32). There was a significant, negative relationship between blood urea nitrogen (BUN) levels and FGF21 (*r* = −0.47, *p* = 0.01) as well as the BUN:Creatinine ratio (*r* = −0.37, *p* = 0.03). Overall, higher levels of FGF21 were associated with decreased glomerular filtration rates (eGFR) (*r* = −0.90, *p* = 0.33). FGF21 was positively associated with bilirubin (*r* = 0.13, *p* = 0.27) and aspartate transaminase (AST) (*r* = 0.22, *p* = 0.14) and negatively associated with the liver enzymes alkaline phosphatase (*r* = −0.03, *p* = 0.44) and alanine transaminase (ALT) (*r* = −0.21, *p* = 0.16). In males, this association with ALT was statistically significant (*r* = −0.53, *p* = 0.04). Relationships between FGF21 and metabolic panel variables are shown in table 3.

**Table 3.** Correlations between FGF21 and lipid panel and metabolic panel variables. * = statistically significant correlation. HDL, high density lipoprotein; LDL, low density lipoprotein; BUN, blood urea nitrogen; eGFR, estimated glomerular filtration rate; AST, aspartate aminotransferase; ALT, alanine aminotransferase.

| Correlate | Whole Group |  | Females |  | Males |  |
| --- | --- | --- | --- | --- | --- | --- |
|  | *r* | *p* | *r* | *p* | *r* | *p* |
| Cholesterol | -0.04 | 0.43 | 0.54 | 0.03* | -0.33 | 0.15 |
| Triglycerides | 0.00 | 0.5 | 0.12 | 0.34 | -0.21 | 0.26 |
| HDL | 0.15 | 0.24 | 0.39 | 0.09 | 0.18 | 0.29 |
| LDL | 0.14 | 0.26 | 0.23 | 0.23 | -0.15 | 0.32 |
| Glucose | 0.10 | 0.32 | 0.24 | 0.21 | 0.10 | 0.37 |
| BUN | -0.47 | 0.01* | -0.32 | 0.15 | -0.58 | 0.03* |
| Creatinine | -0.06 | 0.39 | 0.12 | 0.34 | -0.10 | 0.37 |
| BUN:Creatinine Ratio | -0.37 | 0.03* | -0.20 | 0.25 | -0.56 | 0.03* |
| eGFR | -0.90 | 0.33 | -0.23 | 0.23 | 0.07 | 0.42 |
| Bilirubin | 0.13 | 0.27 | 0.33 | 0.14 | 0.47 | 0.06 |
| Alkaline Phosphatase | -0.03 | 0.44 | 0.35 | 0.12 | -0.01 | 0.49 |
| AST | 0.22 | 0.14 | 0.31 | 0.15 | 0.23 | 0.24 |
| ALT | -0.21 | 0.16 | 0.09 | 0.38 | -0.53 | 0.04* |

### Lipid Panel

Cholesterol was very weakly correlated to FGF21 for the whole group (*r* = −0.04, *p* = 0.43), but in females there was a strong, positive correlation between FGF21 and cholesterol (*r* = 0.54, *p* = 0.03). Both HDL (*r* = 0.15, *p* = 0.24) and LDL (*r* = 0.14, *p* = 0.26) cholesterol were positively related to FGF21 for the whole group. Correlations for the lipid panel results are in table 3.

### Blood Pressure and Heart Rate

Systolic blood pressure (SBP) (*r* = −0.33, *p* = 0.05) had a stronger relationship to serum FGF21 levels than diastolic blood pressure (DBP) (*r* = −0.08, *p* = 0.36). Pulse pressure had a negative correlation with FGF21 (*r* = −0.31, *p* = 0.07) and in females this relationship was statistically significant with a large effect size (*r* = −0.76, *p<*0.01). Resting pulse (RP) had a weak, positive relationship with blood FGF21 levels (*r* = 0.18, *p* = 0.19). Results for the blood pressure and heart rate data are shown in table 4.

**Table 4.** Correlations between FGF21 and cardiovascular variables. * = statistically significant correlation. SBP, systolic blood pressure; DBP, diastolic blood pressure; MAP, mean arterial pressure; PP, pulse pressure; RP, resting pulse; RPP, rate pressure product.

| Correlate | Whole Group |  | Females |  | Males |  |
| --- | --- | --- | --- | --- | --- | --- |
|  | *r* | *p* | *r* | *p* | *r* | *p* |
| SBP | -0.33 | 0.05 | -0.49 | 0.10 | 0.00 | 0.50 |
| DBP | -0.08 | 0.36 | -0.20 | 0.26 | -0.01 | 0.49 |
| MAP | -0.24 | 0.13 | -0.39 | 0.10 | -0.01 | 0.49 |
| PP | -0.31 | 0.07 | -0.76 | <0.01* | 0.01 | 0.49 |
| RP | 0.18 | 0.19 | 0.21 | 0.24 | 0.14 | 0.34 |
| RPP | -0.06 | 0.39 | -0.13 | 0.34 | 0.09 | 0.39 |

## Discussion

In the present study, we demonstrate relationships between serum FGF21 levels and the protein’s upstream and downstream signaling pathways *in vivo*.

We first investigated the relationship between FGF21 and anthropometric variables. In this study, we witnessed a negative relationship between FGF21 and bodyweight, BMI, and waist circumference, this is in contrast to prior studies that have witnessed a positive relationship between FGF21 and obesity both in mice(Fisher *et al*., 2010; Ding *et al*., 2012) and in humans.(Zhang *et al*., 2008; Dushay *et al*., 2010; Roesch *et al*., 2015) The paradoxical relationship of obese individuals having greater levels of circulating FGF21 than normal weight individuals has led to the hypothesis of FGF21 resistance where FGF21 levels are elevated as a compensatory mechanism to obesity, but are unable to exert their normal effect. Our inclusion criteria restricted the BMI to normal weight individuals which may help explain this discrepancy.

Next, we investigated the relationship between FGF21 and daily activity and metabolic variables. Acute exercise studies have consistently shown a brief increase in FGF21 post-exercise,(Kim *et al*., 2013; Hansen *et al*., 2015; Morville *et al*., 2018) while the results of exercise training studies have not been as consistent.(Kruse *et al*., 2017) Previous observation studies have shown a negative relationship between circulating FGF21 and daily physical activity(Cuevas-Ramos *et al*., 2010) along with cardiorespiratory fitness.(Taniguchi *et al*., 2014) Similar results are presented in this study with FGF21 being negatively correlated to VO_2peak_ and average daily activity with a trend for greater effect size as activity intensity increased. Furthermore, we saw a positive relationship between FGF21 and sedentary time providing further evidence that more active individuals have lower circulating levels of FGF21. These findings may relate back to the idea of FGF21 resistance, where more active individuals have a healthier metabolism and do not experience elevated FGF21 levels.

One of the largest effect sizes in our study was in the investigation of sleep and FGF21 levels. FGF21 has been shown to oscillate in a circadian pattern(Yu *et al*., 2011) and sleep plays an important role in maintaining metabolic health(Grandner, 2017) which may underscore the relationship between sleep and FGF21. Interestingly, we observed differences both in the magnitude and direction of the sleep-FGF21 correlation coefficient between males and females. Although our methods for measuring sleep were limited and did not breakdown sleep architecture, prior research has demonstrated sex differences in sleep architecture, with females exhibiting greater amounts of slow wave sleep and males experiencing greater amounts of non-rapid eye-movement sleep.(Mallampalli and Carter, 2014) It is possible that sex difference in sleep architecture could have contributed to this finding. To date, little research has been done on the relationship between FGF21 and sleep, the results of this study indicate it may be an important area for future investigation. In particular, future research could be done to examine the effect of different sleep architectures on FGF21 levels.

One of the promising findings in FGF21 biology from animal models is that exogenous FGF21 administration can induce weight loss. This weight loss has been demonstrated to be the result of an increased energy expenditure coupled with an increase in fat oxidation.(Coskun *et al*., 2008) In agreement with the work done in animals we observed that circulating FGF21 levels were associated with a greater percent of fuel oxidation coming from fatty acids. In contrast, FGF21 was linked with a lower RMR in our study, particularly in females. The participants in our study were all weight-stable, indicating they were at a metabolic homeostasis which may explain this finding. It is also possible that endogenous FGF21 concentrations do not exert strong control over RMR and that higher levels derived through pharmacological administration would be needed to increase RMR. The relationship between metabolic rate and FGF21 in humans is a subject for further investigation, in particular to determine if sex differences observed in this study are replicable in other cohorts.

Transcription of the *FGF21* gene can be promoted by fatty acid and carbohydrate stimuli.(Uebanso *et al*., 2011) We found a negative relationship between FGF21 and fat consumption. In HepG2 cells, fatty acids, particularly unsaturated fatty acids, promoted increased FGF21 secretion(Mai *et al*., 2009) and in mice, fasting and the associated rise in free fatty acids can increase circulating FGF21 levels.(Fazeli *et al*., 2015) However, in humans a much longer fast is required than in mice to induce similar increases in FGF21. Ketogenic diets in humans also have a minimal effect on FGF21 levels, unlike in mice.(Fazeli *et al*., 2015) Furthermore, a high fat diet did not increase human FGF21 levels.(Lundsgaard *et al*., 2016) This study provides further evidence that the link between FGF21 and fatty acids may not be as strong in humans as it is in mice. We found a positive relationship between dietary carbohydrate consumption and FGF21 levels, with particularly strong effect sizes for added sugar and sucrose. This is consistent with other studies that have indicated an acute overconsumption of carbohydrates is associated with a rise in FGF21 levels.(Dushay *et al*., 2015; Lundsgaard *et al*., 2016) Thus, while FGF21 levels may rise following an acute consumption of carbohydrates, our study indicates that higher levels of habitual carbohydrate intake are also associated with greater FGF21 concentrations. Protein does not seem to influence FGF21 to the same extent as other macronutrients, except that low levels of protein consumption are associated with increased FGF21 levels as part of the fasting response.(Laeger *et al*., 2014) Our observation of a negative relationship between protein consumption and FGF21 is consistent with this idea. Finally, nutrients that are metabolized by the liver such as ethanol and fructose have been shown to dramatically increase FGF21 levels in acute feeding studies.(Dushay *et al*., 2015; Søberg *et al*., 2018) We saw a positive relationship between both ethanol and fructose with FGF21 levels. While one-third of our sample reported zero ethanol consumption over the prior month, limiting the strength of association in the present study, the positive association suggests that liver metabolized nutrients upregulate FGF21 in healthy humans.

In our metabolic panel results, we witnessed a smaller correlation between FGF21 and fasting glucose than has been witnessed in other studies;(Cuevas-Ramos *et al*., 2010; Roesch *et al*., 2015) however, our study was limited to metabolically healthy individuals which may explain this difference. We observed a statistically significant, negative relationship between BUN levels and BUN:Creatinine ratios and FGF21. The role of FGF21 in kidney function is an emerging topic of study,(Anuwatmatee *et al*., 2019; Nakano *et al*., 2019) that will require specific intervention studies to fully elucidate.

When exogenous FGF21 has been given to humans, researchers have noted an improvement in dyslipidemia.(Gaich *et al*., 2013) FGF21 is believed to inhibit cholesterol synthesis by inhibiting sterol regulatory element-binding protein-2 (Srebp-2) in the liver.(Jin, Lin and Xu, 2016) In contrast to pharmaceutical intervention studies, we noticed a significant correlation between cholesterol and FGF21 in females and a positive relationship between LDL and FGF21. Circulating FGF21 is believed to derive from hepatic origin; moderate correlations were found between FGF21 and traditional liver markers (AST, ALT, Bilirubin) in this study.

Finally, we explored the relationship between FGF21 and indices of blood pressure. We observed a negative relationship between FGF21 and systolic and diastolic blood pressure. We also witnessed a negative relationship between FGF21 and pulse pressure, indicating that FGF21 is associated with greater arterial compliance. This is in agreement with prior work which found that FGF21 protected against atherosclerosis.(Jin, Lin and Xu, 2016) In one study, a decrease in FGF21 was associated with an increase in arterial compliance;(Yang *et al*., 2011) however, that finding was part of an exercise intervention study, where decreased FGF21 levels may have been caused by weight loss and where improvements of arterial compliance may have arisen from other factors. In summary, we present evidence that there may be a link between FGF21 and arterial compliance, though this is unlikely to result via direct link and may instead be the result of an intermediary such as adiponectin.(Jin, Lin and Xu, 2016)

Our study is not without limitations. First, this was an observational study with a relatively small number of participants. Therefore, we did not find statistical significance in many variables, but by examining the effect size and direction of the relationship we are able to provide meaningful insights into the relationship between FGF21 and other physiological factors. By averaging physiological variables and blood FGF21 concentrations across two separate days we hoped to overcome the common problem of inter-day variation in FGF21 within the same individual.(Gälman *et al*., 2008; Christodoulides *et al*., 2009) We also employed a fairly homogenous, metabolically healthy cohort of individuals. Endogenous FGF21 levels have been shown to be altered in metabolically unhealthy individuals, and as such our results may not be translatable to other populations.

## Conclusion

In this study we present mixed results that suggest normal circulating levels of FGF21 may be related to physiologic and lifestyle factors commonly associated with its action, such as physical activity and carbohydrate consumption, while other factors such as RMR and LDL-cholesterol may not be as related as animal studies and pharmacological interventions indicate. We present strong correlations between FGF21 and sleep and kidney function which are areas prime for future intervention studies.

## Acknowledgements

None

